# Predicting and Analyzing the Response to Selection on Correlated Characters

**DOI:** 10.1101/348896

**Authors:** Tom J.M Van Dooren, Cerisse E. Allen, Patrícia Beldade

## Abstract

The breeder’s equation generally provides robust predictions for the short-term evolution of single characters. When selection targets two or more characters simultaneously, there are often large discrepancies between predicted and observed responses. We assessed how well this standard model predicts responses to bivariate selection on wing color pattern characteristics in the tropical butterfly *Bicyclus anynana*. In separate laboratory selection experiments, two sets of serially repeated eyespots were subjected to ten generations of concerted and antagonistic selection for either size or color composition. We compared predicted and actual selection responses over successive generations, using the phenotypic data, selection differentials, and estimates of the genetic variance-covariance matrix ***G***. We found differences in the precision of predictions between directions of selection but did not find any evidence of systematic biases in our predictions depending on the direction of selection. Our investigation revealed significant environmental effects on trait evolution across generations. When these were accounted for, predictions using the standard model improved considerably. In the experiment on eyespot size, secondary splitting of selection lines allowed the estimation of changes in ***G*** after nine generations of selection. Changes were not in general agreement with expectations from the breeder’s equation. A contour plot of prediction errors across trait space suggests that directional epistasis in the eyespot genotype-phenotype map might occur but estimates of changes in ***G*** are too model-dependent to verify whether they agree with that hypothesis. Altogether, our results underscore the need for quantitative genetics to investigate and estimate potential effects of multivariate non-linear genotype-phenotype maps and of environmental effects on ***G***.

## Introduction

The ‘breeder’s equation’, which we also call the ‘infinitesimal model’ (Barton et al. 2017; Turelli 2017) is a robust and useful tool for understanding evolutionary dynamics. While simulations suggest that deviations from the Gaussian distributions assumed in this model usually have small effects (Turelli & Barton 1994; Zhang & Hill 2005), observed selection responses can differ from infinitesimal predictions (Sheridan 1988; Hill 2010; Roff 2007). When two or more characters are selected simultaneously, predictions for the multivariate response are often much less accurate (Falconer & Mackay 1996; Roff 2007). Multiple studies suggest that the response becomes more difficult to predict when two characters are selected in opposing directions (antagonistic selection) compared with selection in the same direction (concerted selection; Bell & Burris 1973; Falconer & Mackay 1996 1996; Roff 2007). Several alternative explanations for this poor predictability of antagonistic selection have been proposed (reviewed in Roff 2007), as well as methods to assess them. A first possibility is that the breeder’s equation would predict approximately correct, but that the data analysis is more cumbersome. Secondly, there might simply be too few examples comparing *a priori* predictions with empirical results to draw robust conclusions about the predictability of bivariate or multivariate evolution. Another possibility could be that predicting the response of many traits might require a much more elaborate model selection procedure and increased risk of prediction biases. Alternatively, the standard quantitative genetic models may be deemed inadequate - either too simplified, or failing to account for critical underlying factors, such as developmental interactions, that might limit phenotypic evolution (Pigliucci & Schlichtling 1997) in a way not covered by the equations. In data analysis, when we are unable to predict responses well, explanations will therefore range from inference issues to a fundamental failure to capture properties of the biological system well. Failing at predicting responses to selection almost appears more exciting than obtaining good predictions. It is, however, possible to embed the breeder’s equation into a model selection and model simplification framework from which potentially improved predictions and much insight can be gained when mechanistic models are used (Le Rouzic et al. 2011). Environmental effects and changes to the genotype-phenotype map which have been invoked to explain results of selection experiments (Okada & Hardin 1967) can be fitted to the data. However the approach is currently only available for single traits and can therefore not be used yet to understand differences in performance of the breeder’s equation in a multivariate setting.

Some model simplifications leading to inaccurate predictions might not be core assumptions of the breeder’s equation and would therefore not warrant rejecting it, when they rather follow from common practise and usually remain untested. For instance, short-term changes in additive genetic variance and covariance due to selection over a few generations are often assumed to be negligible, and are typically ignored. However, the effects of selection and drift can change these parameters within a relatively small number of generations. Ignoring such changes might be one cause of poor predictability of selection response in a number of analyses. Short-term selection experiments (≥ 5 generations; Hill 2011) are useful for assessing changes in components of ***G*** during the course of selection (Hill 2011; Heath et al 1995; Martinez et al. 2000; Meyer & Hill 1991; Beniwal et al. 1992). Predicting multivariate selection-induced effects on ***G*** (the genetic variance-covariance matrix) remains involved, despite a great deal of theory (e.g., Lande 1979; Barton & Turelli 1987; Johnson & Barton 2005) and empirical work (e.g., Meyer & Hill 1991; Beniwal et al. 1992; Heath et al 1995).

There are several ways to test whether the standard multivariate breeder’s equation is appropriate in a given context. Demonstrating non-Gaussian genotype and phenotype distributions may invalidate the model, but not immediately demonstrate that selection responses are poorly predicted. In non-pedigreed populations subject to artificial laboratory selection, standard selection analysis uses least-squares techniques to assess model fit (Falconer & Mackay 1996). By fitting the standard model to observed responses in a series of different selection lines, patterns in the residuals of the predicted means can be investigated, in a strategy that is frequently used for model validation in other contexts. The usefulness of this exercise relies on accurately estimating both ***G*** and ***β*** (the selection gradient), which will depend on the design of the artificial selection experiment. When the experimental design allows ***G*** to be estimated separately both at the start and end of the experiment, it is possible to determine whether ***G*** estimated after several generations of selection still fits the predictions of the infinitesimal model (the “Gaussian population” approximation, Turelli 2017), given the starting estimate of ***G*** and the empirical selection gradient. Thus, it is possible to test whether ***G*** has changed during the course of selection, and whether such changes are predicted by the infinitesimal model. Because the assumptions of the standard infinitesimal model are likely violated after many generations of selection during which ***G*** may undergo substantial changes, tests of the infinitesimal model are most appropriately applied to selection experiments with small to intermediate numbers of generations, where the model is generally believed to perform well. When changes in ***G*** are estimated, it is far from straightforward to assign multivariate estimated changes in different treatment groups (Arnold et al. 2008) to mechanisms. As stated already, statistical modelling tools to do that in the context of time series of selection responses are not immediately available. It can also happen that the infinitesimal model does produce adequate predictions, but that ***G*** has changed in a way not anticipated by it.

To assess how well the standard infinitesimal model predicts bivariate evolution and to investigate whether this model still improves our understanding of the biological system, we analyzed phenotypic data from two artificial selection experiments targeting correlated eyespot characters in the tropical butterfly *Bicyclus anynana* (Nymphalidae: Satyrinae). These characters were 1) eyespot size (relative to wing size), a trait largely determined by the strength of the eyespot-organizing morphogen produced by the cells at the center of the presumptive eyespot, and 2) eyespot color-composition (proportion of black and gold), a trait probably determined by the sensitivity thresholds to an eyespot-inducing signal (see Beldade & Brakefield 2002, Beldade et al. 2008, Allen et al. 2008). We currently lack direct evidence concerning the number of loci or distributions of allelic effects underlying these eyespot characteristics in *B. anynana*. A significant portion of the standing variation for size and color composition appears to be additive (Monteiro et al. 1994; Monteiro et al. 1997; Beldade et al. 2002b), and allelic variation at the Distal-less locus accounts for up to 20% of the difference between lines selected for the size of either the anterior dorsal forewing eyespot EyeA or the posterior dorsal forewing eyespot EyeP (Beldade et al. 2002a). Very little is known about the genetic architecture underlying eyespot color composition, though models suggest that the diffusion gradient-threshold mechanisms employed in eyespot development likely generate nonlinear gene effects (Gilchrist & Nijhout 2001).

In each experiment, pairs of eyespots were selected in both concerted and antagonistic directions (analyses of the phenotypic responses are reported by Allen et al. 2008; Beldade et al. 2002b). The structure of the ***G*** matrix seemed comparable between size and color composition traits in previous analysis (Allen et al. 2008), such that different outcomes between selection experiments prompted a discussion on the relevance of quantitative genetic methods for this model system.

We re-analyzed the data more elaborately than before and with ***G*** re-estimated for each experiment. To predict selection responses, we used different estimates of ***G*** per experiment: one estimate obtained from a separate breeding experiment using the unselected stock population, and another estimate obtained using data from the base population (prior to selection) and the first generation after selection. Descendants of the base population were partitioned into several lines selected in several directions, targeting eyespot size (in the first experiment; Beldade et al. 2002b,c) or eyespot color composition (in the second experiment; Allen et al. 2008). Model selection and comparison allowed us to determine whether the choice of model effects fitted biased our estimates of ***G***. Using those estimates of ***G***, we subsequently predicted selection responses and assessed model fit to the selection data by analyzing the residuals from these predictions. To avoid analyzing spurious patterns in these data resulting from a sub-optimal fit, we made a further effort to select and fit a model which best predicted the actual selection response and minimized the overall variance of residuals. In addition, the experimental design of the eyespot size experiment allowed us to estimate ***G*** again after nine generations of selection and compare that estimate with infinitesimal model predictions.

## Materials and Methods

### Artificial selection experiments

In two separate experiments, we selected for either the relative size of two dorsal forewing eyespots (EyeA and EyeP, for the anterior and posterior eyespots, respectively, typically found on the forewing), or the color composition of two ventral hindwing eyespots (Eye4 and Eye6, for the fourth and sixth eyespots, respectively, of the seven typically found on that wing surface) of *Bicyclus anynana* (Figure 1). The starting population (Gen0, for generation zero) for each experiment was derived from the same outbred stock maintained in the laboratory for > 100 generations at high *N*_e_ (Brakefield et al. 2001). In both experiments, only females were selected and selection was maintained at similar intensities for 10 generations. Per line and per generation, we measured 150–200 females for size (mean ± SE: 173 ± 3) and 140–240 females for color composition (mean ± SE: 209 ± 5). We selected 40 females per line every generation; in the size experiment this number decreased to 35 females per line between generations 5–10. Details including choice of traits for selection, selection criteria, selection procedure, and analysis of the rates of response to selection are described in (Allen et al. 2008; Beldade et al. 2002 b,c). Here we report eyespot size (relative to wing size) and color composition (size of the black disc relative to total eyespot size) as percentages.

**Figure 1.**
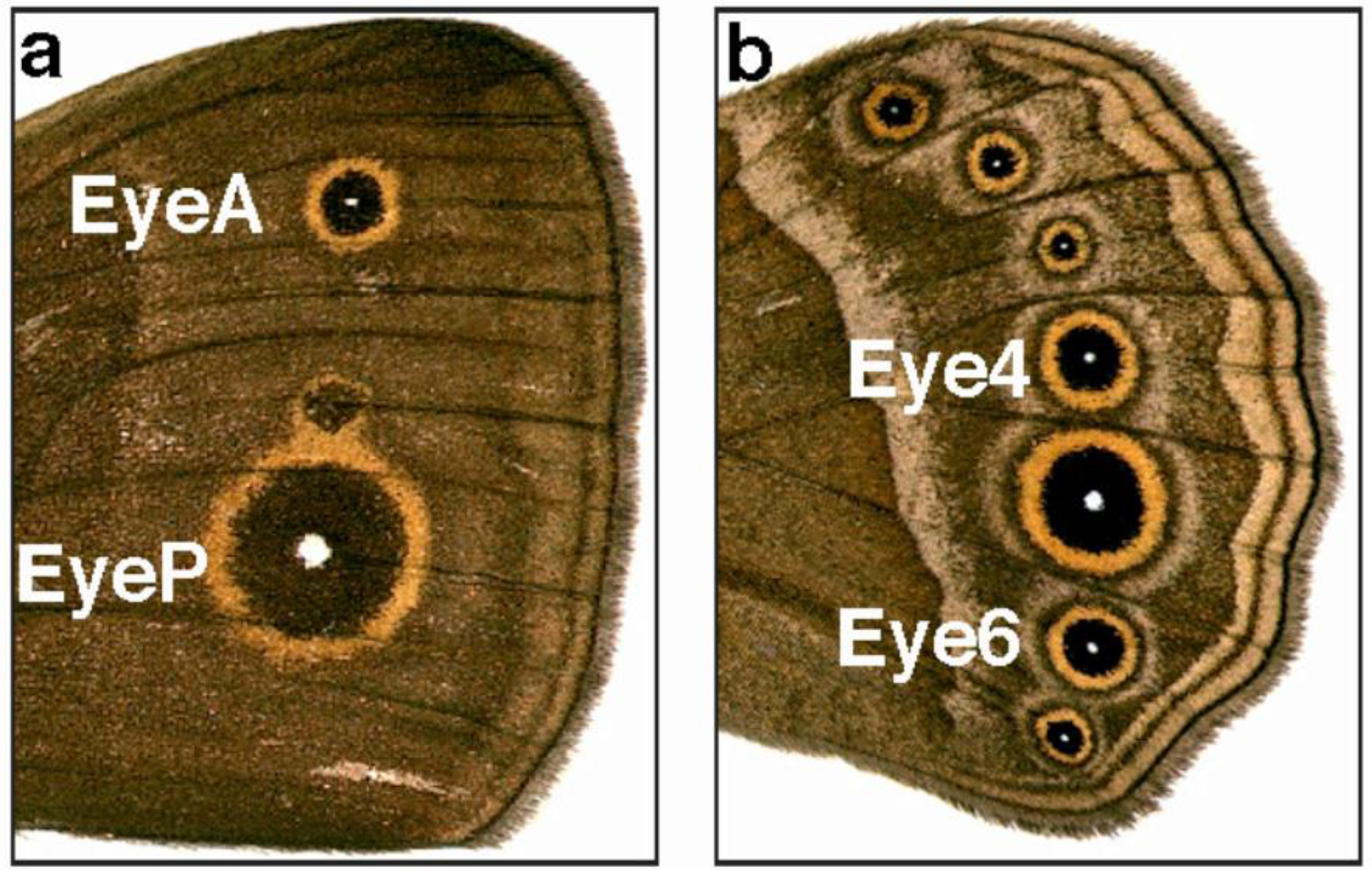
Location and form of the dorsal forewing and ventral hindwing eyespots of *Bicyclus anynana*. The dorsal forewing in **A** illustrates the locations of the anterior (EyeA) and posterior (EyeP) eyespots. Selection targeted combinations of eyespot sizes; size was measured relative to total wing size (see text). The ventral hindwing in **B** illustrates all seven eyespots, with markers indicating the locations of the two eyespots (Eye4 and Eye6) targeted by simultaneous selection for color composition. Color composition was estimated as the diameter of the inner black ring relative to total eyespot diameter (see text for details).

In both experiments, we established three types of lines from the starting population (see Fig. 2): 1) antagonistic selection lines where two eyespots were selected in opposite directions (e.g. larger EyeA and smaller EyeP), orthogonal to the main axis of phenotypic and genetic correlations among eyespots; 2) concerted selection lines where two eyespots were selected in the same direction (e.g. larger EyeA and EyeP), parallel to the main axis of phenotypic and genetic correlations among eyespots; and, 3) unselected control (UC) lines. Each direction of selection was replicated twice. In both experiments, lines were selected for 10 generations, but as several of the color lines were lost through error in the final generation, response is shown for that experiment after nine generations of selection only.

**Figure 2.**
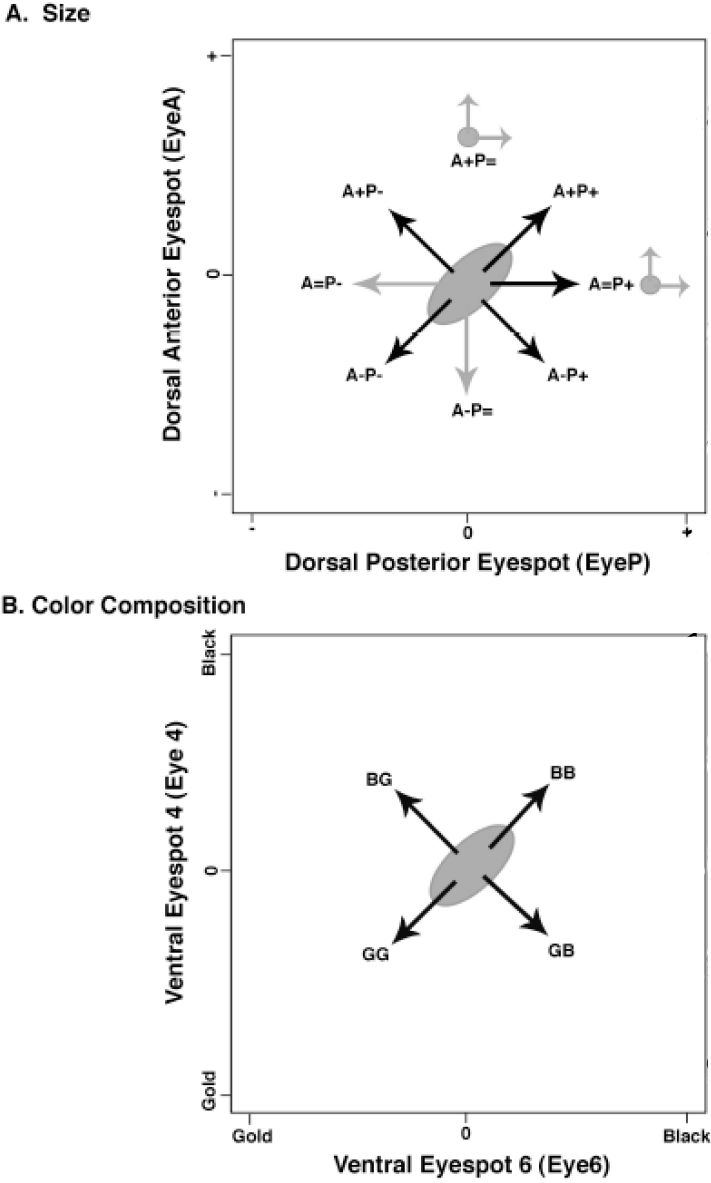
Directions of artificial selection imposed on eyespot size and eyespot color composition in *Bicyclus anynana*. Ellipses represent the location of the starting (stock) population. Selection occurred for ten generations (black arrows) in most directions; gray arrows signify directions where only a single generation of selection occurred. **A**, artificial selection simultaneously targeted the size (relative to wing size) of the anterior (EyeA) and posterior (EyeP) eyespots on the dorsal forewing surface. Eyespots were selected for increased size (+), decreased size (−), or constant size (=). After nine generations (small gray circles), butterflies from lines A+P= and A=P+ were split into subpopulations and selected along two orthogonal directions for one generation (short gray arrows). **B**, artificial selection simultaneously targeted the color composition (amount of black relative to total size) of the fourth and sixth eyespots on the ventral hindwing. Eyespots Eye4 and Eye6 were selected for either increased proportion of gold (G) or increased proportion of black (B) scales, for ten generations. Selection on eyespot color composition occurred only along the concerted (both eyespots selected in the same direction) or antagonistic (each eyespot selected in a different direction) axes, and there was no further splitting of lines.

In the eyespot size experiment, two additional types of lines were established from the starting population (Fig. 2): 4) uncoupling selection lines where one eyespot was subjected to directional selection and the other eyespot was simultaneously subjected to stabilizing selection (e.g. larger EyeA and constant EyeP); and 5) re-split lines where, after nine generations of selection, two of these directional/stabilizing selection lines were each split into two populations and selected either along the original axis or an orthogonal axis for an additional generation.

### Estimates of G

#### Outbred laboratory stock population

We used a paternal half-sib breeding design {Lynch & Walsh 1998} to estimate additive genetic variance and covariance of four eyespot characters in our stock population at a time between the two selection experiments. We randomly selected 100 virgin males from the outbred stock at adult eclosion and allowed each male to mate sequentially with at least two virgin females. At hatching, ~30 eggs per female were transferred to mesh rearing cages and fed on young maize plants ad libitum until pupation. Full-sib offspring were reared together but densities were kept low to minimize interactions and competition between individuals. Rearing cages were moved every four days to randomize environmental effects within the growth chamber. Emerging adult offspring were allowed several hours for their wings to expand and fully harden before being frozen for later measurements.

Five female offspring were randomly selected from each of 174 full-sib families (representing 87 sires who successfully produced offspring by two dams each) and dorsal forewing eyespots EyeA and EyeP, and ventral hindwing eyespots Eye4 and Eye6 were measured as described on the left wings only of each individual. We used our nested breeding design to obtain REML estimates of sire, dam (nested within sires), and progeny variance and covariance components in the software package ASReml (VSN International, 2006). We tested for differences between the dam and sire genetic covariance matrices using a likelihood ratio test, which is a conservative approach (Pinheiro & Bates 2000; Verbeke & Molenberghs 2000). Both the dam and sire covariance matrices are reported.

#### *G_0_* in the starting population

With equal phenotypic and genetic variances in both sexes, random mating among parents, and no environmental changes, the expected bivariate mean phenotypic trait vector in the first generation after artificial selection, for a selection line *i*, is described by

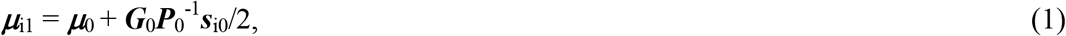

Where ***μ***_0_ is the mean phenotypic trait vector in the base/starting population, ***s***_i0_ is the selection differential for females from the base population that initiate line *i*, ***G***_0_ the genetic variance-covariance matrix, and ***P***_0_ the phenotypic variance-covariance matrix in the base population.

When there are common environmental effects on mean trait values in generation one, these can be added as vector ***μ***_e1_ to the right-hand side of Eqn. (1). When the phenotypic trait values are multivariate normal, then for each individual in line *i* of generation *j* (0 or 1), the probability density function of the individual trait vector ***x*** (which contains two trait values for each experiment) is

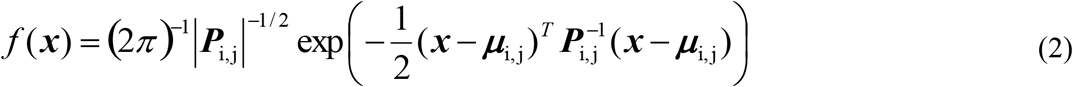

To calculate the likelihood of a dataset with observations in the base population and generation one, given a set of parameter values, the *f*(***x***) of all individuals in the dataset must be multiplied. Equation (2) can be used to model the dependence of trait values in generation one on trait values in generation zero by replacing ***μ***_i1_ by Eqn. (1). This is a regression model. Using maximum likelihood techniques, we can then estimate the bivariate means ***μ***_i,0_ and the variance components of ***P***_i,j_ per line per generation, and estimate ***G***_0_ and ***P***_0_ in the base population. The selection differentials act as observed covariates, and are not estimated in the ML model.

We compared models where common environmental effects, ***μ***_e1_, were included or excluded, and where ***P*** remained fixed or was allowed to vary between selection lines or directions of selection. Since both the means and variances of the bivariate normal distribution can differ between these models, we used maximum likelihood instead of restricted maximum likelihood estimation of parameters (Verbeke & Molenberghs 1997). All ML fitting was done using R statistical software (Ihaka & Gentleman 1996). Although a disadvantage of ML is that the phenotypic variance estimates are biased downward, this estimation is asymptotically efficient (Cox & Hinkley 1974). We obtained ML estimates of all model parameters and their approximate confidence intervals, based on the curvature of the likelihood function, or, for parameter estimates very near to the boundary of the parameter space, by direct profile likelihood intervals (Pawitan, 2001).

We used likelihood ratio tests to compare nested models. Since these tests are not available for non-nested models, we could only compare the AIC (Aikake Information Criterion, Akaike 1973) between them. This kind of model comparison is not frequentist inference. As the AIC, we report twice the negative log-likelihood plus the number of parameters in the model. In model comparison, the model with the smallest AIC value is preferred. To ensure positive estimates of phenotypic variances, we used a log link function for parameterization. We checked normality assumptions by inspecting normal probability plots of residuals from the most parameter-rich models we fitted.

#### Least squares estimates of *G*

The half-sib estimate of ***G*** and the ML estimate of ***G***_0_ do not necessarily minimize the difference between actual and predicted response. To find the ***G*** matrix minimizing the summed squared differences between predicted and actual selection response, we conducted a minimization routine assuming fixed ***P*** and ***G*** across generations and no environmental effects. As a measure of model fit, we calculated differences between predicted and actual response per line *i* and generation *j*, Σ_i,j_ and then summed all the squared differences, Σ_i,j_^T^Σ_i,j_ across all selection lines (not including controls) and generations. This measure is a ‘residual sum of squares’ and we determined the ***G*** minimizing it. This analysis did not include the four size lines that were selected in a new direction after the split at Gen9. Unlike the ML estimate of ***G***_0_ (or the half-sib estimate), this least-squares estimate depends on trait values in all generations.

### Predicted versus actual responses to selection

#### Performance of different estimators

After estimating the genetic variance-covariance matrix in three different ways, we used these estimates to predict selection responses in all subsequent generations and compared the fit of different models to the observed data. Eqn. (1) is easily extended to predict the response from one generation to the next by replacing the generation index 0 by *j* and 1 by *j + 1*.

As a starting point, we modeled selection response assuming: a) that ***G*** did not change during each experiment (separate models incorporated either the ML estimate of ***G***_0_, the half-sib estimates or the LS estimate); b) ***P*** remained unchanged and identical to starting population values during the experiment; and c), no generation-specific environmental effects on mean trait values. The LS estimate of ***G*** necessarily had to perform best among these models.

We attempted to improve model fit by modelling changes in ***P*** and incorporating these in the predictions. The time- and line-dependent models we investigated assumed multivariate normality and used ML estimation of means and (co)variances (Eqn. 2). However, modeling phenotypic covariance matrices and using the resulting parameter estimates to predict selection response increased the residual sum of squares (i.e., reduced model fit). Simply substituting sample estimates of ***P*** in Eqn. (1) also reduced model fit. Thus, the ‘best fit’ model for ***P*** used in subsequent steps was actually the one where ***P*** was fixed at the estimate of the starting population (Gen0).

#### Accounting for the effects of selection on *G*

Under the infinitesimal model, genetic variance components change due to gametic-phase linkage disequilibrium (Bulmer 1971; Lynch & Walsh 1998). The expected change in the genetic variance-covariance matrix for selection line *i*, in generation *j* after selection, is

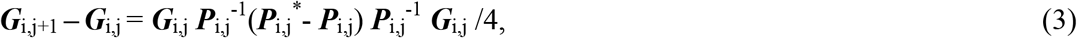

where ***P***_i,j_* is the phenotypic variance among the individuals of line *i* in generation *j* selected to contribute to the next generation. The expected change in the genetic variance-covariance matrix due to this type of linkage disequilibrium was calculated for each selection line *i* and following each generation *j* of selection, and incorporated into equations to predict the selection response. We also modeled the ***P***_i,j_* and found that predictions were best when we used the sample statistics per generation. We compared the subsequent fit with the previous models where ***G*** was assumed fixed.

#### Variation in predictability and bias

Because accuracy of predicted responses may differ among lines and traits, we checked for line-specific or direction-specific systematic differences between predicted and actual selection responses (bias) and for line- or direction-specific changes in the variances of these differences (predictability). We also checked for the presence of common environmental effects on generation means, which are thus generation-specific biases.

We calculated differences between predicted and actual response, Σ_i,j_, based on the best fit model achieved so far. To investigate line- or direction-specific bias and common environmental effects, we fitted repeated measurements models (Lindsey 1999) to the differences, Σ_i,j_. We used the elliptic function of Lindsey’s growth library (Lindsey 1999) with auto-correlated errors, normally distributed residuals, and changes in the variance of Σ_i,j_ between lines, and fit models to each trait separately. First, models included all effects (line-specific, direction-specific, common environment per generation) and were later simplified using backward model selection by means of likelihood ratio tests. Because autocorrelation was weak, we subsequently fit generalized least squares models using Venables and Ripley’s (2002) GLS function and did model selection on these. A dependence of predictability on the direction of selection, as expected from other studies, was assessed by testing whether the variances of the differences varied significantly between lines or directions of selection, biases were investigated by testing whether certain averages differed significantly from zero. We examined the residual sums of squares again to determine to which extent models incorporating these line, direction or generation effects improved our ability to predict selection responses (i.e., reduced residual sum of squares). These analyses included all selection and control lines.

#### Environmental effects

It is customary to adjust selection responses with environmental effects estimated from control lines only. For that reason, we also estimated environmental effects using control lines alone and checked how much these reduced the residual sums of squares when they were incorporated in predictions of selection response.

### Changes in G between generations zero and nine of the size selection experiment

To determine whether selection on eyespot size altered ***G*** between Gen0 and Gen9, we applied the same approach for estimating ***G***_0_ (above) to estimate ***G*** in Gen9, using the four size selection lines that were re-split at Gen9 (the lines at Gen9 constitute the new ‘starting population’, and each line has two descendant lines following selection; see Fig. 2). Trait values for the base population (Gen0), Gen1, a given (split) population in Gen9 and its offspring in Gen10 were combined in a single model fit. This allowed us to directly estimate the differences between components of ***G*** in the base population and a given descendant population in Gen9.

For each selection line, we fitted a model that allowed changes in all three parameters of ***G*** and sub-models that allowed from none to two parameters of ***G*** to change. The models for each line also included global environmental effects on character means estimated for Gen10 and line-specific changes in the phenotypic variance between Gen9-Gen10 (see above). We again used likelihood ratio tests to compare nested models with different numbers of parameters, and the AIC to compare models with equal numbers of parameters (e.g., to compare two models that each included one parameter change in ***G***). In this way, we selected a model that best described the changes in ***G*** for the empirical data; these changes were not constrained to follow the patterns predicted by the infinitesimal model (Eqn. 3). We then compared our estimated changes in G between Gen0-Gen9 with predicted changes according the infinitesimal model (Eqn. 3).

As a final investigation of the potential changes in ***G***, we followed-up on a suggestion detailed by Le Rouzic et al. (2011) that local acceleration or deceleration of selection responses unexplained by the breeder’s equation can be caused by changes in the local curvature of the genotype-phenotype map. We thus fitted bivariate generalized additive models (gam, Wood 2017) to prediction errors per trait remaining when environmental effects are accounted for. This is different from our analysis of predictability and bias in that we don’t test for differences in bias between lines but for local bias variation across trait space. For gam’s where thin plate regression splines of average anterior and posterior eyespot size in the population had significant effects on the prediction error, we made contour plots of the pattern of model predictions to see whether the starting population and the populations with secondary splittings were situated at trait values close to contours with positive (augmented response) or negative (lagging response) values. If that is the case, directional epistasis might cause ***G*** to change.

## Results

### Estimates of G

#### Outbred laboratory stock population

Substantial additive genetic variances (V_A_), covariances, and genetic correlations (*r*_G_) for eyespot size and color composition were detected in the unselected stock population using paternal half-sib analysis (Table 1). Because observed sire variances were consistently larger than dam variances (Table 1) we report both estimates separately and make separate predictions using sire and dam genetic variance (see below).

**Table 1.** Estimates of the variance-covariance matrix ***G*** used to predict selection response.

| Population | Size |  |  | Color composition |  |  | Method |
| --- | --- | --- | --- | --- | --- | --- | --- |
| | $V_A$ | $V_A$ | $r_G$ | $V_A$ | $V_A$ | $r_G$ | |
|  | EyeA | EyeP |  | Eye4 | Eye6 |  |  |
| **Unselected Stock (sire)** | 20.7 (6.4) | 55.1 (14.7) | 0.69 (0.13) | 9.7 (4.1) | 15.1 (4.0) | 0.71 (0.15) | REML |
| **Unselected stock (dam)** | 15.4 (5.3) | 30.4 (10.6) | 0.41 (0.24) | 6.8 (4.3) | 4.1 (3.0) | 0.78 (0.34) | REML |
| **Gen0** | 17.3 (1.0) | 30.0 (1.5) | 0.56 (0.03) | 5.4 (0.8) | 4.4 (0.7) | 1.00<br>(0.83 - 1.00) | ML |
| **Gen0 (constant)** | 17.4 (0.6) | 29.0 (1.9) | 0.57 (0.02) | 4.9 (0.6) | 3.8 (0.4) | 0.78 (0.07) | LS |
| **Gen0 (updated)** | 19.3 (0.7) | 37.1 (2.8) | 0.72 (0.03) | 5.8 (0.8) | 4.7 (0.6) | 0.81 (0.06) | LS |

The estimates obtained from the unselected stock also included (four) pairwise genetic correlations between eyespot size and color composition traits, which are not reported here.

However, the standard error of half-sib estimates are relatively large compared to estimates obtained from the selection data (see below), and the sire and dam variances were not significantly different in a likelihood ratio test (χ^2^ = 6.80, *df* = 10, *p* = 0.744).

#### *G*_0_ in the starting population

The details of ML model selection and parameter predictions for Gen0 are given in Table 2 (eyespot size) and Table 3 (eyespot color composition). The best fit model for both data sets included global (common to all lines) environmental effects which changed phenotype means between Gen0 and Gen1 (Tables 2 and 3): After one generation of selection, there was a positive environmental effects on the size of eyespots EyeA and EyeP (Table 2; Mean environmental effect), and a negative environmental effect on the relative blackness of eyespot Eye4 (Table 3; Mean environmental effect). For the eyespot size dataset, the best-fit model incorporated changes in phenotypic variances and covariances that were specific to each direction of selection (but without any obvious pattern of change related to the direction; Table 2). Model fit was poorer (higher AIC values) when models incorporated either line-specific changes in P or differentiated between groups of antagonistic and concerted selection lines. For the color composition dataset, in contrast, the best fit model incorporated a global change in P across all lines between Gen0 and Gen1 (Table 3).

**Table 2.** Maximum likelihood parameter estimates for the starting population and first offspring generation in the eyespot size selection experiment.

| Eyespot Size | ML Model I | ML Model II | ML Model III |
| --- | --- | --- | --- |
| AIC | 47448 | 47230 | **47184** |
| Number of Parameters | 11 | 13 | **40** |
| Generation 0 |  |  |  |
| Mean EyeA | 26.3 (0.1) | 26.3 (0.1) | **26.3 (0.1)** |
| Mean EyeP | 57.3 (0.1) | 57.3 (0.1) | **57.3 (0.1)** |
| $V_G$ EyeA | 17.8 (1.0) | 17.0 (1.0) | **17.3 (1.0)** |
| $V_G$ EyeP | 30.6 (1.6) | 30.4 (1.5) | **30.0 (1.5)** |
| $r_G$ (EyeA, EyeP) | 0.57 (0.03) | 0.57 (0.03) | **0.56 (0.03)** |
| Generation 1 |  |  |  |
| Mean Environmental effect EyeA | 0 | 2.2 (0.1) | **2.2 (0.1)** |
| Mean Environmental effect EyeP | 0 | 1.4 (0.2) | **1.5 (0.2)** |
| Modeled changes in phenotypic variances | Shared among all lines | Shared among all lines | Per direction of selection |

Estimates of character means, genetic variances ( $V_G$ ), genetic correlations ( $r_G$ ), and their standard errors (in parenthesis) are given for Generation 0, the starting population before selection. Estimates are given for each of three ML models: Model I, the model with the fewest parameters, no common environmental effect, and all lines share a common change in phenotypic variance between Gen0 and Gen1; Model II, the model with the fewest parameters plus a common environmental effect; Model III is the model with the lowest AIC among all models. It allows changes in phenotypic variances between generations. Estimates from the overall best fit model are in bold.

**Table 3.**
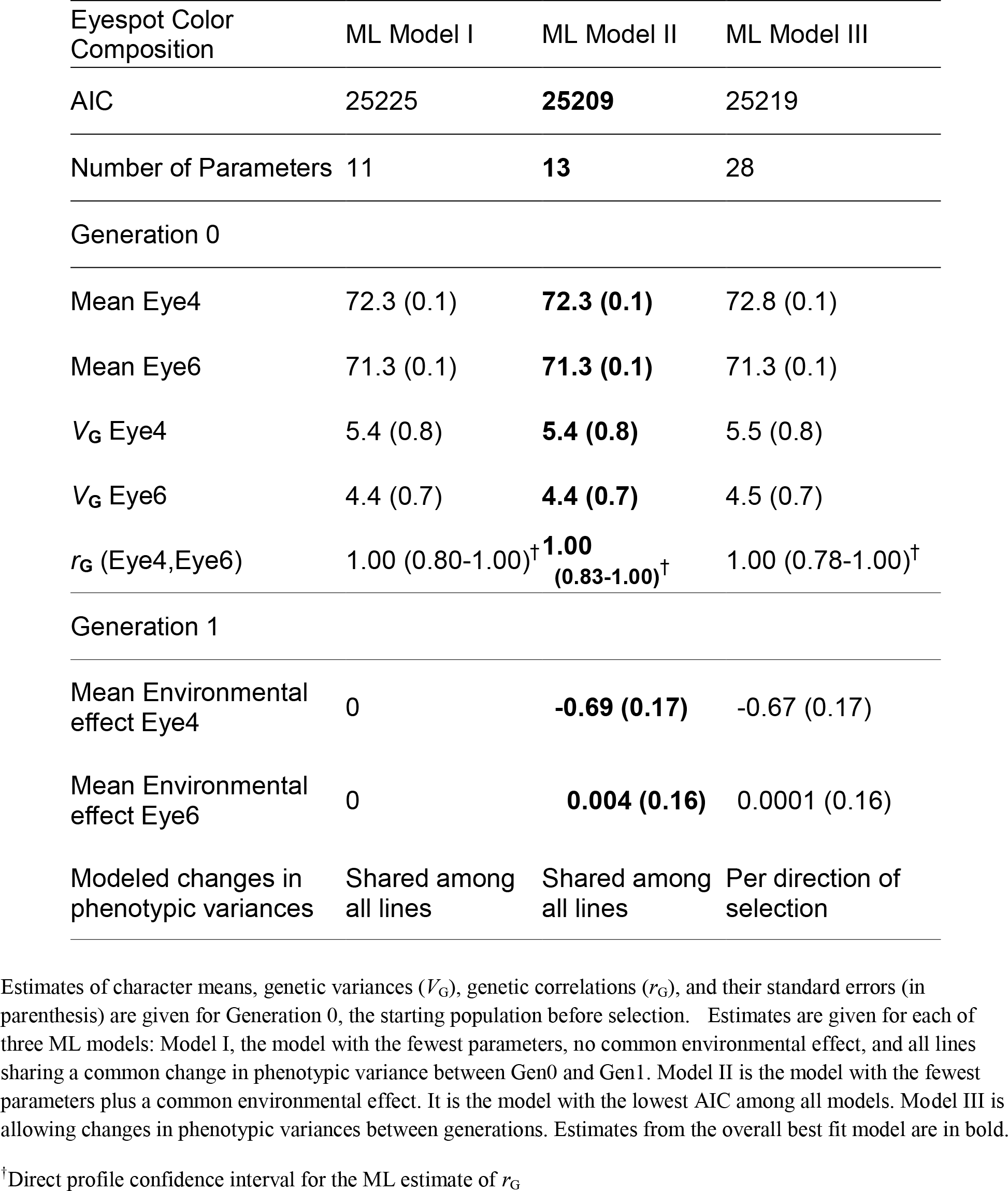
Maximum likelihood parameter estimates for the starting population and first offspring generation in the eyespot color composition selection experiment.

| Eyespot Color Composition | ML Model I | ML Model II | ML Model III |
| --- | --- | --- | --- |
| AIC | 25225 | **25209** | 25219 |
| Number of Parameters | 11 | **13** | 28 |
| Generation 0 |  |  |  |
| Mean Eye4 | 72.3 (0.1) | **72.3 (0.1)** | 72.8 (0.1) |
| Mean Eye6 | 71.3 (0.1) | **71.3 (0.1)** | 71.3 (0.1) |
| $V_G$ Eye4 | 5.4 (0.8) | **5.4 (0.8)** | 5.5 (0.8) |
| $V_G$ Eye6 | 4.4 (0.7) | **4.4 (0.7)** | 4.5 (0.7) |
| $r_G$ (Eye4, Eye6) | 1.00 (0.80-1.00)<sup>†</sup> | **1.00 (0.83-1.00)<sup>†</sup>** | 1.00 (0.78-1.00)<sup>†</sup> |
| Generation 1 |  |  |  |
| Mean Environmental effect Eye4 | 0 | **-0.69 (0.17)** | -0.67 (0.17) |
| Mean Environmental effect Eye6 | 0 | **0.004 (0.16)** | 0.0001 (0.16) |
| Modeled changes in phenotypic variances | Shared among all lines | Shared among all lines | Per direction of selection |

Estimates of character means, genetic variances ( $V_G$ ), genetic correlations ( $r_G$ ), and their standard errors (in parenthesis) are given for Generation 0, the starting population before selection. Estimates are given for each of three ML models: Model I, the model with the fewest parameters, no common environmental effect, and all lines sharing a common change in phenotypic variance between Gen0 and Gen1. Model II is the model with the fewest parameters plus a common environmental effect. It is the model with the lowest AIC among all models. Model III is allowing changes in phenotypic variances between generations. Estimates from the overall best fit model are in bold.
<sup>†</sup>Direct profile confidence interval for the ML estimate of $r_G$

The ML genetic parameter estimates for Gen0 were similar across all models (Tables 2 & 3: see parameter estimates for ML models I, II, and III); thus, model selection did not appear to bias estimates. For the best fit models, the estimated genetic correlation between eyespots EyeA and EyeP = 0.56 ± 0.02, and the estimated genetic correlation between eyespots Eye4 and Eye6 = 1.0 (profile likelihood confidence interval = 0.83 - 1.00).

#### Least squares estimates

For each dataset, we calculated a least-squares (realized) estimate of ***G*** (Table 1) that minimized the summed squared differences between predicted and actual selection response, across all generations. In general, the least-squares estimates are concordant with ML estimates and fall within their range of expected error (compare to estimates in Tables 2 & 3); however, the estimate for the genetic correlation for color composition of Eye4 and Eye6 (*r*_G_ = 0.78 ± 0.07) is slightly lower than the ML estimate.

### Predicted versus observed responses to selection

#### Performance of different estimators

Table 4 shows the sums of squared differences (residual sum of squares; RSS) between predicted and actual responses to selection under a number of different model conditions. As a measure of global model fit (all lines, all generations, per dataset), we compared these residual sums of squares to the total sum of squared differences between the actual responses per line per generation and the overall mean. First we held ***G*** and ***P*** constant and did not include environmental effects. The sire (REML) estimate of ***G*** in the stock population produced the largest mismatch between predicted and observed responses (the largest RSS, Table 4): 3.4% of the total sums of squares for the eyespot size data, and 17.4% of the total for the color composition data. The dam (REML) estimate of ***G*** substantially increased model fit relative to the sire estimate in both experiments (Table 4). Both the ML estimate of ***G***_0_ and the LS estimate (realized ***G***) produced slight improvements over the dam estimate (Table 4, both datasets). The smallest residual sum of squares for eyespot size under the basic conditions is 1.6% of the total (LS estimate; Table 4); it is 6.4% of the total for eyespot color composition (LS estimate; Table 4).

**Table 4.** Unexplained variation in the predicted response to selection.

| Residual sum of squares | Total sum of squares | **G** | Environmental effects |
| --- | --- | --- | --- |
| Eyespot size | 29059 |  |  |
| 523.66 |  | Fixed; REML (dam) | Not included |
| 986.35 |  | Fixed; REML (sire) | Not included |
| 491.23 |  | Fixed; ML (**G**<sub>0</sub>) | Not included |
| 490.18 |  | Fixed; LS | Not included |
| 602.47 |  | Updated; ML (**G**<sub>0</sub>) | Not included |
| 571.06 |  | Updated; LS | Not included |
| 289.31 |  | Fixed; LS | Included |
| 388.81 |  | Updated; LS | Included |
| Eyespot color composition | 1287 |  |  |
| 85.85 |  | Fixed; REML (dam) | Not included |
| 224.10 |  | Fixed; REML (sire) | Not included |
| 86.65 |  | Fixed; ML (**G**<sub>0</sub>) | Not included |
| 82.65 |  | Fixed; LS | Not included |
| 83.29 |  | Updated; ML (**G**<sub>0</sub>) | Not included |
| 81.82 |  | Updated; LS | Not included |
| 39.30 |  | Fixed; LS | Included |
| 38.02 |  | Updated; LS | Included |

#### Accounting for the effects of selection on *G*

In general, adjusting ***G*** each generation to account for linkage disequilibrium generated by selection (according to Eqn. 3) did not affect model fit relative to constant ***G***. For the size dataset, the residual sums of squares increased when ***G*** was allowed to change across generations (Table 4: LS updated, RSS = 571.06; LS fixed, RSS = 490.18). For the color composition dataset, accounting for changes in ***G*** due to linkage disequilibrium slightly improved model fit but only reduced RSS by ~1% relative to models with constant ***G*** (Table 4: LS updated, RSS = 81.82; LS fixed, RSS = 82.65).

#### Variation in predictability and bias

We found that the predictability of actual selection responses differed between individual selection lines in both datasets. Predictability of selection response also differed between eyespots within an experiment. In the size dataset, predictability of the selection response of EyeA varied significantly between directions of selection (heterogeneity of error variances: LRT = 43.62, *df* = 14, *p* < 0.001), but this could not be simplified by grouping concerted and antagonistic directions (LRT = 18.78, *df* = 2, *p* < 0.001). Predictability of EyeA response appeared to vary between different directions of selection: lines in the A-P+ direction had the largest error variances and the A+P− and A+P+ lines (see Figure 3) had the smallest error variance (<10% of the largest line-specific error variance). Overall, predictability of the response of EyeP was not significantly different between selection lines (LRT = 19.18, *df* = 14, *p* = 0.16). We therefore did not find any evidence that as a group, predictability differed between concerted and antagonistic selection lines in this experiment.

**Figure 3.**
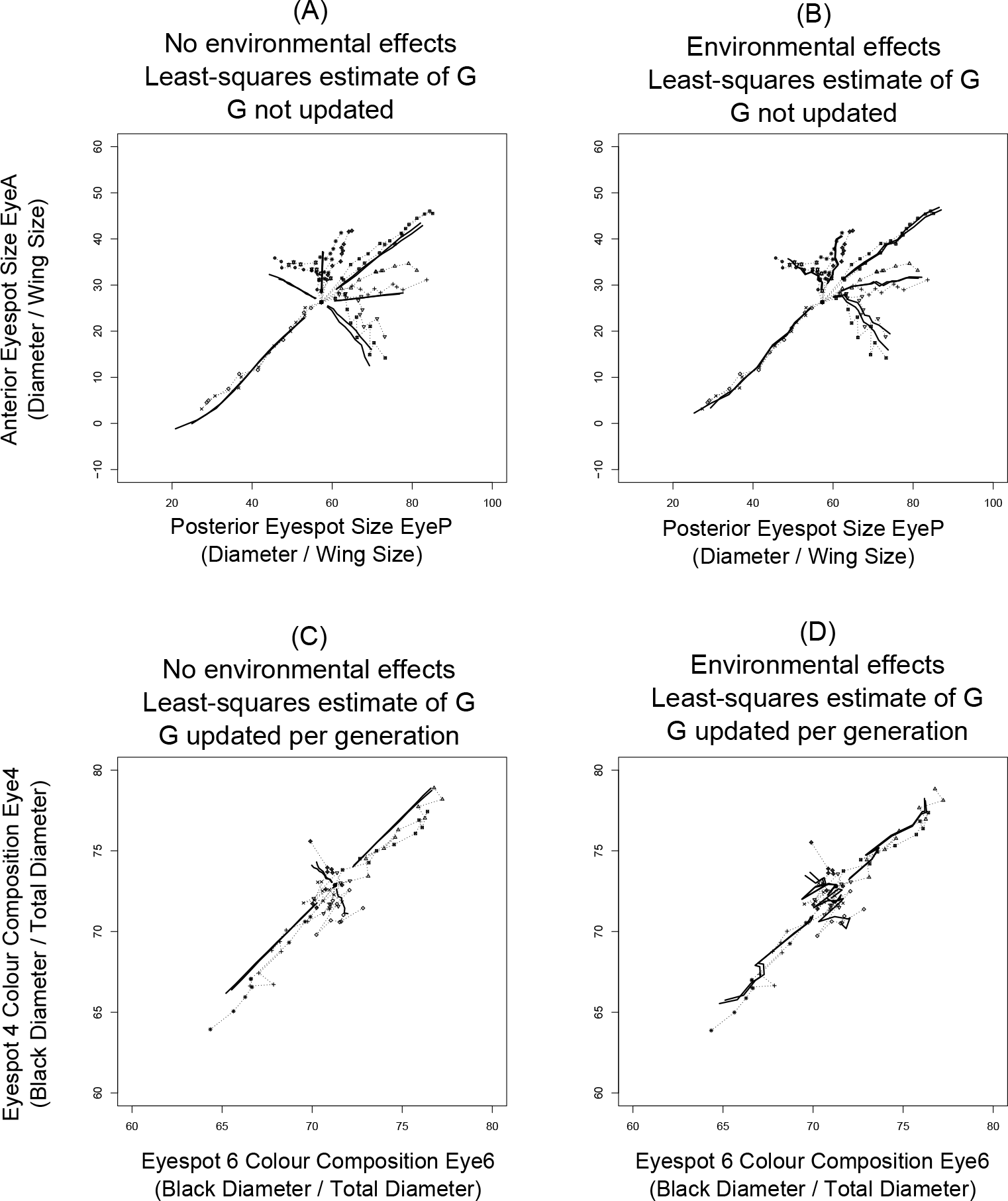
Including environmental effects improves the correspondence between predicted and observed responses to artificial selection. Predicted responses (solid lines) and observed selection responses (dashed lines connecting data points) are shown for each replicate in each selected direction. For each experiment, we show the best fit model without environmental effects included (**A** and **C**), and the best fit model with environmental effects included (**B** and **D**). For eyespot size (**A** and **B**), ***G*** is fixed at starting values. For eyespot color composition, the least-squares estimate of ***G*** was updated each generation to account for the effects of selection on genetic variances and covariances. See Table 4 for details of the model fitting and choice of the best fit model in each experiment.

In the color composition dataset, predictability of the selection response varied significantly between lines for Eye6 (LRT = 18.52, *df* = 9, *p* = 0.03) but not Eye4 (LRT = 11.38, *df* = 9, *p* = 0.25). The predictability of selection responses did not vary between selection directions, and did not differ between concerted and antagonistic selection lines. Concerted lines 4B6B_2_, and 4G6G_2_ and the antagonistic line 4G6B_2_ (see Figure 2), had the smallest error variance (each line with < 20% of the largest line-specific error variance) and the concerted line 4B6B_1_ had the largest error variance. Despite the fact that individual lines differed in the predictability of selection response, the per-line average errors were not significantly different from zero in either the size or color experiment. This means that responses were never consistently over- or underestimated in any of the selection lines or selection directions and there were no significant line biases in the responses of concerted versus antagonistic lines.

#### Environmental effects

We found significant common environmental effects on eyespot phenotype means within generations and across all lines for both datasets (all *p* < 0.0001 Table 5). Environment affected the mean size of EyeA and EyeP, and the mean color composition of Eye4 and Eye6 independent of the effects of selection (also see above and Tables 2 and 3).

**Table 5.** Global environmental effects estimated for the eyespot size and color composition selection experiments.

| SIZE |  |  |
| --- | --- | --- |
| Generation | EyeA Environmental effect | EyeP Environmental effect |
| 1 | **2.10 (1.70 - 2.50)** | **1.31 (0.70 - 1.93)** |
| 2 | **0.93 (0.53 - 1.33)** | **1.17 (0.56 - 1.79)** |
| 3 | 0.02 (-0.38 - 0.42) | 0.12 (-0.49 - 0.74) |
| 4 | 0.19 (-0.21 - 0.59) | **1.20 (0.59 - 1.82)** |
| 5 | 0.29 (-0.11 - 0.69) | **-0.72 (-1.33 - -0.11)** |
| 6 | **-1.09 (-1.49 - -0.69)** | -0.10 (-0.71 - 0.51) |
| 7 | **0.74 (0.34 - 1.14)** | 0.07 (-0.55 - 0.68) |
| 8 | 0.19 (-0.59 - 0.21) | 0.11 (-0.73 - 0.50) |
| 9 | **0.66 (0.26 - 1.07)** | 0.29 (-0.33 - 0.90) |
| 10 | -0.25 (-0.65 - 0.15) | **1.21 (0.60 - 1.83)** |
| Color composition | Eye4 Environmental effect | Eye6 Environmental effect |
| 1 | **-0.72 (-1.18 - -0.26)** | -0.28 (-0.64 - 0.09) |
| 2 | 0.45 (-0.01 - 0.91) | **0.57 (0.20 - 0.93)** |
| 3 | -0.02 (-0.48 - 0.44) | -0.25 (-0.62 - 0.11) |
| 4 | **-1.13 (-1.59 - -0.66)** | **-1.34 (-1.71 - -0.98)** |
| 5 | **0.57 (0.11 - 1.04)** | **0.84 (0.48 - 1.20)** |
| 6 | 0.06 (-0.41 - 0.52) | **0.67 (0.31 - 1.04)** |
| 7 | 0.26 (-0.20 - 0.72) | 0.03 (-0.33 - 0.40) |
| 8 | -0.37 (-0.83 - 0.10) | -0.19 (-0.55 - 0.18) |
| 9 | 0.32 (-0.14 - 0.78) | **-0.46 (-0.83 - -0.10)** |

Global environmental effects estimated across all lines, per generation, for eyespot size and eyespot color composition experiments. Environmental effects on each eyespot were estimated using all lines, including unselected controls (see Methods). Estimates are shown with their 95% confidence intervals in parentheses. Values in bold indicate environmental effects for which the confidence interval does not include zero.

Incorporating common environmental effects into our predictions for selection responses in each dataset visibly improved the correspondence between predicted and actual response (Figs. 3d, 4d) and substantially reduced the residual sums of squares experiments (size: to 289.31, a 41% reduction; color composition: to 38.02, a 54% reduction; Table 4).

When we used only the unselected control (UC) lines to estimate the between-line, within-generation environmental effects, the only significant global environmental effects were for EyeA in Gen1 and Gen6. This method did not reveal any significant common environmental effects on EyeP, Eye4, or Eye6 in any generation. Despite this, incorporating environmental effects estimated from the UC lines still substantially improved the correspondence between predicted and actual responses to selection (sum of squared differences = 359.21, and 55.00 for eyespot size and color composition, respectively; Table 4) relative to models that did not incorporate environmental effects.

### Changes in G between generations zero and nine of the size selection experiment

A subset of the size selection lines was used to estimate changes in genetic variances and covariances between Gen0 and Gen9. We detected significant changes in ***G*** in two of the stabilizing-directional lines, and a trend in a third line (Table 6). In these three lines, the best fit model (lowest AIC) included a change in at least one parameter of ***G***. However, the parameter estimates themselves are highly model dependent. Estimates change drastically depending on the particular model (Table 6). Therefore individual estimates must be interpreted with caution.

**Table 6.** Predicted changes in ***G*** according to the infinitesimal model, versus changes in ***G*** estimated from eyespot size selection lines after nine generations of selection.

| Selection line | Parameter | ML Model (i)<br>Estimated change<br>(S.E.) | ML Model (ii)<br>Estimated change<br>(S.E.) | ML Model (iii)<br>Estimated change<br>(S.E.) | Infinitesimal<br>model<br>Predicted<br>change |
| --- | --- | --- | --- | --- | --- |
| A= P+<sub>(1)</sub> | V<sub>A</sub> EyeA | 7.20 (2.81) | 0 | 0 | -8.63 (0.59) |
|  | V<sub>A</sub> EyeP | -9.53 (4.33) | -13.60 (4.35) | 0 | -17.33 (2.45) |
| | $r_G$ (EyeA,<br>EyeP) | 0.03 (0.07) | 0 | 0 | -0.10 (0.004) |
|  | AIC | 50436.52 | 50433.70 | 50437.26 |  |
| | | | $P = 0.02$ | | |
| A= P+<sub>(2)</sub> | V<sub>A</sub> EyeA | -2.33 (4.37) | 54.89 (5.10) | 0 | -5.14 (0.56) |
|  | V<sub>A</sub> EyeP | -1.54 (4.11) | 21.80 (4.66) | 0 | -17.30 (2.47) |
| | $r_G$ (EyeA,<br>EyeP) | -0.66 (0.14) | 0 | 0 | -0.17 (0.01) |
|  | AIC | 50723.16 | 50721.14 | 50730.20 |  |
| | | | $P < 0.001$ | | |
| A+ P=<sub>(1)</sub> | V<sub>A</sub> EyeA | 4.54 (2.57) | 0 | 0 | -10.02 (0.63) |
|  | V<sub>A</sub> EyeP | -2.19 (5.14) | -15.22 (5.02) | 0 | -14.69 (2.09) |
| | $r_G$ (EyeA,<br>EyeP) | 0.19 (0.06) | 0 | 0 | -0.09 (0.01) |
|  | AIC | 50569.18 | 50566.12 | 50567.38 |  |
| | | | $P = 0.07$ | | |
| A+ P=<sub>(2)</sub> | V<sub>A</sub> EyeA | -4.99 (2.91) | 0 | 0 | -9.91 (0.62) |
|  | V<sub>A</sub> EyeP | 1.24 (4.22) | 0 | 0 | -15.96 (2.26) |
| | $r_G$ (EyeA,<br>EyeP) | 0.10 (0.08) | 0 | 0 | -0.08 (0.01) |
|  | AIC | 50063.54 | 50061.30 | 50061.30 |  |
| | | | $P = 1.00$ | | |

Changes in $G$ between the base population (Gen0) and Gen9 were estimated from the subset of size selection lines which were split at Gen9 and their descendants after one additional generation of selection. ML models either allowed all components of $G$ to change, or constrained successive components of $G$ to no change between Gen0-Gen9. Estimates (with their standard errors) and the associated AIC are given for three models: (i) the model with unconstrained $G$ ; (ii) the model with minimum AIC among all models fitted; (iii) the model with all changes constrained to zero. Predicted changes in $G$ between Gen0-Gen9 according to the infinitesimal model, accounting for linkage disequilibrium, were calculated according to Eqn (3). P values are for likelihood ratio tests comparing the model without any change (iii) with the model with minimum AIC among all models fitted (ii). The standard deviation of the infinitesimal model predicted change was obtained as follows. The matrix $G$ was re-estimated for each combination of lines which still included all selection directions. The predicted change was recalculated for each estimated matrix. The standard deviation of each prediction over the combinations is given.

Regardless of model, in all cases where we detected significant changes in ***G***, the estimated change differed from predicted change (according to Eqn. 3) by at least one standard error (Table 6).

We also applied Eqn. 3 to all lines in both datasets to predict changes in ***G*** due to selection-induced linkage disequilibrium between Gen0-Gen9 (Table 7). We found that the smallest changes due to linkage disequilibrium were predicted in the two stabilizing-directional selection lines (eyespot size) which were re-split at Gen9 (Table 6), and in antagonistic selection lines for size and color composition. In both datasets, Eqn. 3 predicted the largest changes in ***G*** for concerted selection lines. Unfortunately, stabilizing-directional lines (with the smallest predicted changes) were the only lines re-split at Gen9 and available to test actual changes in ***G***.

**Table 7.**
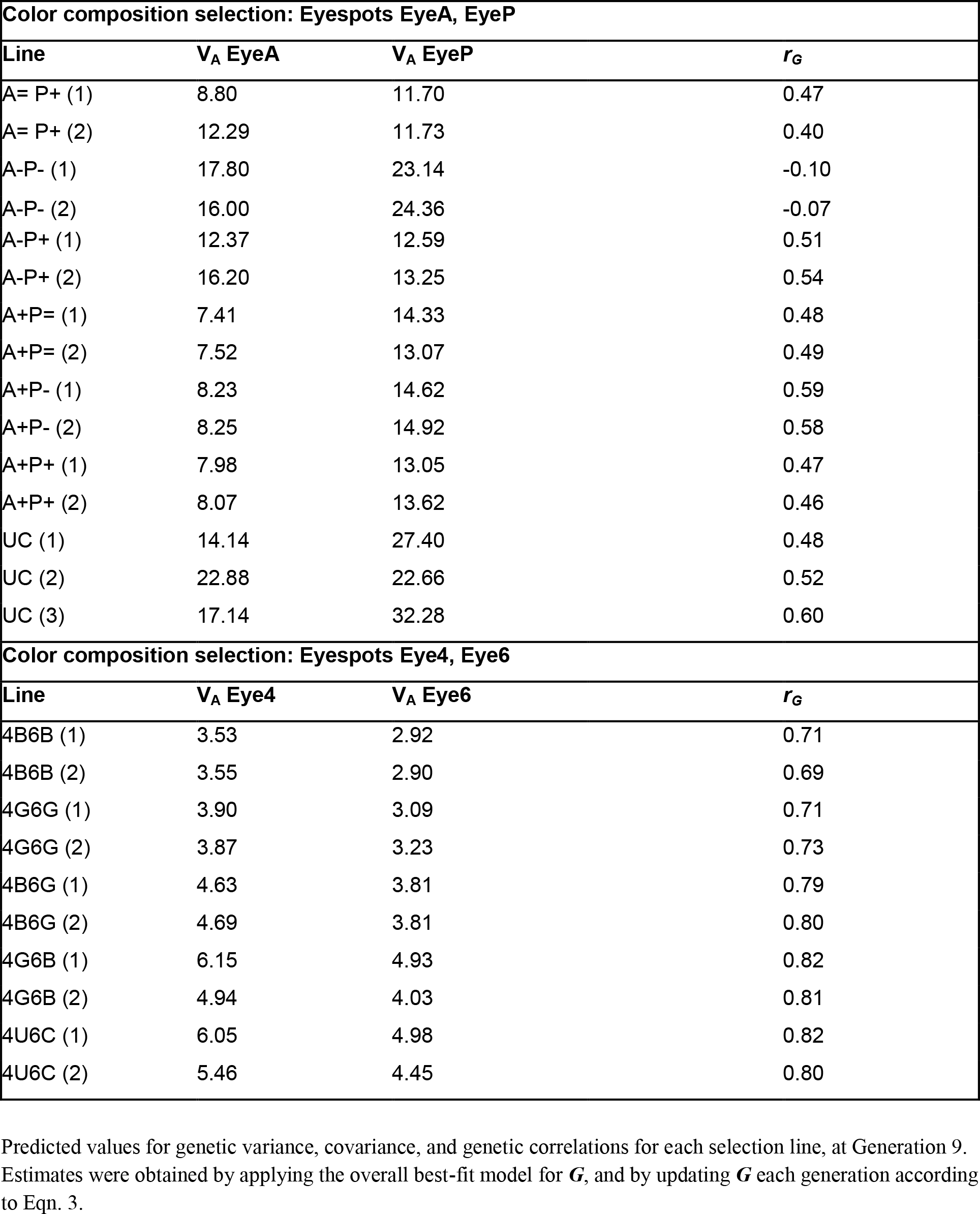
Infinitesimal predictions for genetic parameters at Generation 9 of the eyespot size and color composition experiments.

Even while we could not demonstrate significant bias differences between selection lines, generalized additive models of prediction errors for anterior and posterior eyespot size depended significantly on average values of these two traits in the population (eyeA approximate significance of smooth terms *F*_16.7, 21.4_ = 1.756, *p* = 0.028; eyeP: *F*_13.4, 17.7_ = 2.489, *p* = 0.0015). Figure 4 shows that the local bias is relatively small, which can explain that we did not detect it when selection lines were analyzed as categorical variables. Assuming that directional epistasis causes the pattern, the contour plots of these models (Figure 4) suggest that genotypic trait variances increase with trait means. In comparison with the ancestral population, for example, A=P+ populations then have an increased genetic variance for eyeP.

**Figure 4.**
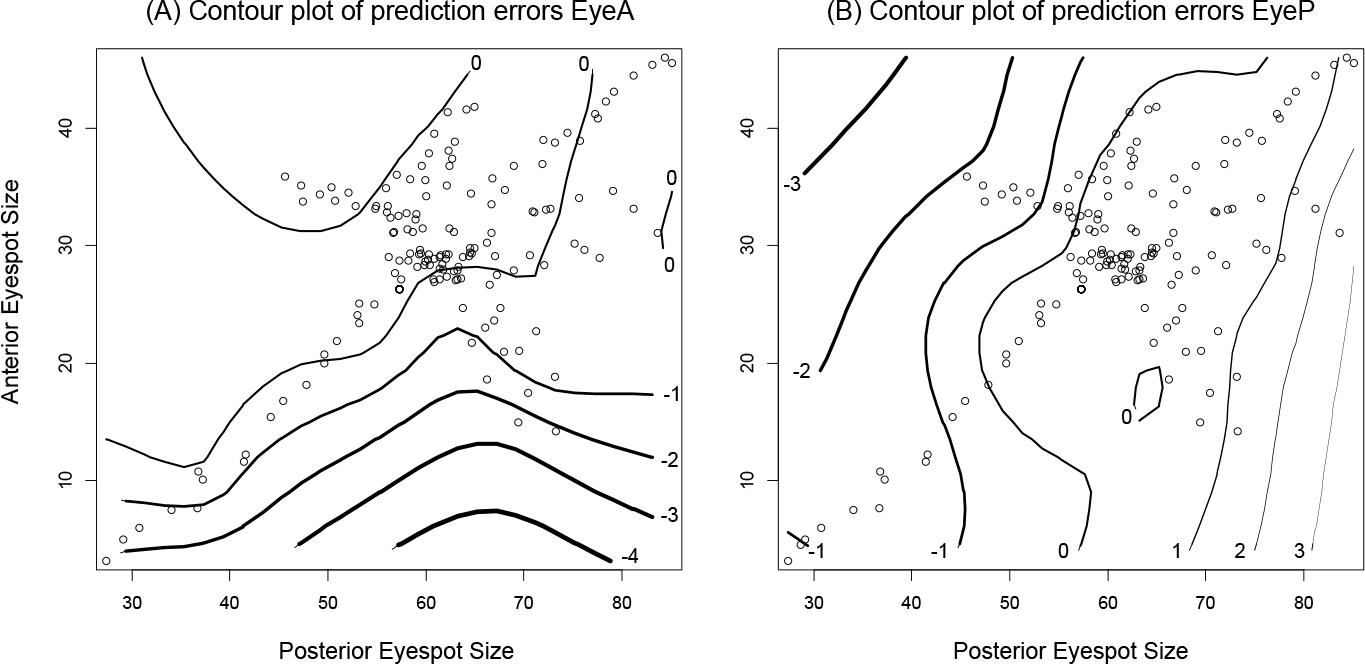
Contour plots of prediction errors in anterior (**A**) and posterior (**B**) eyespot sizes, when shared environmental effects are accounted for. For each trait, the prediction error increases on average with the value of the trait. Such an asymmetric pattern suggests that directional epistasis in the genotype-phenotype map could be present for both traits, and that the effect is that the trait variance positively depends on trait value.

## Discussion

We were able to evaluate the predictive power of the breeder’s equation for two correlated traits using two large artificial selection experiments targeting correlated sets of eyespot characters in *B. anynana* butterflies. Although the standard infinitesimal model predicted evolutionary responses with reasonable accuracy, predictability varied between size and color composition, between individual selection lines within an experiment, and between individual eyespots. Accounting for selection-induced changes in ***G*** (due to linkage disequilibrium, Bulmer 1971) had little, if any, effect on the accuracy of our predictions. Instead we found that accounting for common environmental effects on eyespot phenotypes that were independent of selection significantly improved the correspondence between predicted and observed evolutionary changes in eyespot size and color composition. Using a subset of the data, we detected significant changes in parameters of ***G*** after nine generations of artificial selection. These were not in agreement with predictions from the breeder’s equation.

### Analyzing selection responses in non-pedigreed populations

When pedigree information is available, restricted maximum likelihood (REML) analysis combined with mixed-model analysis of all phenotypic data is used to estimate ***G***_0_ in the base (starting) population (Sorensen and Kennedy 1984). Lacking a pedigree during the selection experiment, we were still able to estimate ***G***_0_ in two different ways. First we used REML mixed model analysis of a half-sib breeding design to estimate ***G*** in the unselected stock population.

Second, we constructed ML models to estimate ***G***_0_, the additive genetic variance-covariance matrix in the starting population prior to selection, for both the eyespot size and color composition experiments. Inspecting sums of squares of the residuals of predicted selection response allowed us to assess whether the stock population estimate of ***G*** (dam estimate) or ***G***_0_ better predicted selection responses across all generations. We found that ***G***_0_ performed better than the stock population estimate, though both estimates provided reasonably accurate predictions for the magnitude and direction of selection response in each experiment.

We used the same ML approach to re-estimate ***G*** later in the experiment, using four size selection lines that were split into sub-lines at Generation 9 and subject to an additional generation of selection. These data were used to investigate whether significant changes in ***G*** occurred during the course of the selection experiment. Although we detected significant changes in ***G*** with this method, parameter estimates of the changes were much less reliable than our initial ML estimation of ***G***_0_ in the starting population. For the starting population, confidence intervals of all estimates were relatively narrow and the estimates themselves were not affected by model selection bias (Tables 2 and 3). In contrast, we found strong model selection bias in our Gen9 estimates of ***G***. Thus it appears that the ML approach we used works well when many lines are started from a single ancestral population and selected in many different directions- the situation that occurred at the onset of both experiments. Our approach is less robust when a small number of lines are started from an ancestral line and selection proceeds in a few limited directions (which occurred during Gen9). An additional disadvantage of our method is that the assumptions of the breeder’s equation may not be satisfied after many generations of selection, when ***G*** is expected to change substantially through changes in allele frequencies (Turelli & Barton 1994). However, that disadvantage applies to all approaches involving the breeder’s equation and does not specifically distinguish our procedure. Simulations are needed to fully assess the power and precision of ML estimates of ***G*** and their dependence on the design of selection experiments. However, our approach is advantageous in that it is not computationally demanding, and in addition, in that the experimental design allowed direct tests for changes in ***G*** across generations not assuming any particular mechanism.

### Does the breeder’s equation predict bivariate responses to selection accurately?

The standard model adequately predicts the direction of evolutionary change for both eyespot size and color composition. This result is perhaps unsurprising- the standard model appears generally robust, even when infinitesimal assumptions are violated (Turelli & Barton 1994; Zhang and Hill 2005). The concordance we found between three separate estimates for ***G*** (REML dam estimate of the unselected stock population; ML estimate for the base population in each experiment; and the LS estimate across all generations of selection), and that fact that all three produced reasonable predictions for short term change (excepting the REML sire estimates; Table 1) suggests that infinitesimal assumptions are reasonable for both datasets. Although estimates of ***G*** from an unselected stock population are generally preferred over realized estimates (Juga & Thompson 1989), our analyses show that both the ML base population estimate and the LS (realized) estimate performed well, while the REML stock population estimate provided less accurate predictions.

Despite considerable unexplained variation in selection response in both experiments, there was no systematic effect of selection direction on predictability- both antagonistic and concerted selection were similarly predictable in each dataset. Some previous work comparing the predictability of antagonistic and concerted selection suggests that short-term, bivariate selection is poorly predicted by the standard model (Berger & Harvey 1975; Bell & Burris 1973). Sheridan & Barker (1974) found that responses in all directions were well predicted during the short-term, but that predictability declined after 22 generations of selection and that changes in genetic correlations did not match expectations. Selection-induced changes in the joint distributions of traits may violate the standard assumption of multivariate normality (Barton & Turelli 1989) and also result in a gradually decreasing predictability of response to selection.

Although selection direction did not influence predictability, predictability of individual characters did vary overall, the relative amount of unexplained variation was substantially different for eyespot size and eyespot color composition, with size being more predictable than color. Within experiments, responses of EyeP and Eye4 were better predicted overall than responses of EyeA and Eye6. In each case, the eyespot with the smaller initial mean value and estimate of V_A_ (EyeA; Eye6) showed significant among-line heterogeneity in the agreement between predicted and observed selection responses. In contrast, there was no significant among-line heterogeneity for the eyespot with the larger mean and estimate of V_A_ (EyeP; Eye4). Whether this suggests an important pattern or follows from a deviation from model assumptions which causes a dependence between trait means and variances requires further analysis.

Two of our attempts to improve the fit of models to the selection response had little effect: accounting for predictable changes in ***G*** caused by selection-induced linkage disequilibrium (the Bulmer effect) had only minimal effects on the residual sums of squares (amount of unexplained variation in response). Similarly, accounting for changes in ***P*** across generations also had no effect. There are many potential sources of variation in predictability of response, including drift, differing allele frequency changes in different lines, nonadditive genetic variation (e.g., directional epistasis), gene-by-environment (GxE) interaction, selection acting on correlated traits, and environmental variation (Falconer & Mackay 1996) or environmental effects on the ***G*** matrix (Wood & Brodie 2015). In our analysis, the most obvious explanation for the overall difference between experiments in amounts of unexplained variation (residual sums of squares after fitting selection response) is sampling effects on the average phenotype in finite populations (Lande, 1976). Sampling variance is proportional to the magnitude of the standing genetic variance; this is in agreement with the larger residual sum of squares for the eyespot size experiment (V_A eyespot size_ > V_A eyespot color_). Though drift is a likely cause of variation in the average trait values each generation (Hill, 1971), we cannot clearly attribute observed changes in ***G*** to drift.

#### Environmental effects are critical for accurate predictions

Global environmental effects (those effects on eyespot character means shared across all lines within a given generation) account for a large proportion of the mismatch between predicted and observed selection responses in both data sets. In both experiments, accounting for environmental variation in the model improved predictions compared with models that accounted for selection-induced changes in ***G*** or ***P*** (Figs. 3 and 4). In contrast with typical analyses that rely on unselected control lines, we used all selected and unselected lines to estimate environmental effects (Falconer & Mackay 1996). Though our approach inevitably leads to a better model fit, it also allows much more accurate estimation of the environmental effect than a comparison with a single control line (Sorensen & Kennedy 1984). This method is probably most robust when selection occurs in different directions with equal numbers of opposing lines, because systematic estimation bias might occur if selection were performed in only a limited number of directions.

Across-generation environmental effects were erratic, without significant trends over time. In addition, significant effects on eyespot means were frequently limited to a single eyespot out of the pair targeted by selection. Fluctuating food-plant quality over the course of each experiment is a possible source of such environmental variation. Food-plant quality and larval crowding can affect many aspects of larval growth and impact both wing pigment production and the appearance of individual wing color pattern characters (Gibbs & Breuker 2006; Talloen et al. 2004). Other aspects of the general rearing environment, such as temperature or humidity, could also have fluctuated during the course of the two experiments and affected particular characters or individual eyespots (Brakefield et al., 1996).

Regardless of their source, environmental effects impact character means, and can push the selection response in a direction opposite to that otherwise predicted (compare panels a, c with b, d in Fig. 3). In Fig. 3, this is particularly clear in antagonistic selection directions, which showed strongly ‘jagged’ responses. Since jaggedness only appears in the predicted trajectories when we include environmental effects in the models, we can clearly identify environment as a major cause of apparent visual irregularity in antagonistic selection responses. General environmental effects can have wide-ranging impacts on many other aspects of the evolutionary process. When genotypes differ in their sensitivity to environmental variables, selection may directly alter environmental variance (Kaufman 1977; Scharloo 1972); changes in environmental sensitivity during an experiment can lead to apparent failure to respond to particular selection pressures (Jinks et al. 1977). In *B. anynana*, substantial family-by-environment variation for wing color pattern characters (Windig 1994) could also account for portions of the variation in selection response that remains after general environmental effects are accounted for.

### Does G change during the course of short-term selection?

By generation nine of the size selection experiment, we observed significant changes to parameters of ***G*** in two of the four lines sampled (and marginally significant changes in a third line; Table 6). In each case, the best fit model indicated that the genetic correlation between EyeA and EyeP remained stable, despite significant changes in V_A_ for one or both of the eyespots. These observed changes in ***G*** are striking when compared with infinitesimal predictions based on Eqn. 3: observed changes in variance components are much larger (and frequently differ in sign) than predicted changes due to selection-induced equilibrium alone. It is possible that our choice of populations that were split again, simply lacked power to detect the full range of changes in ***G***, since infinitesimal predictions led us to expect modest shifts in the magnitude of the genetic correlation *r*_***G***_ in all four of the lines analyzed (Table 5). Estimates of changes depended a lot on whether the some parameters were constrained to zero or not. That suggests that model selection bias is probable and that we should not overinterpret these results. Nevertheless, our analyses suggest that we must consider factors other than drift and gametic-phase disequilibrium to explain the changes observed between Gen0 and Gen9, as these would both produce clear decreases in additive genetic variances. Given the results of the generalized additive models (gam’s) fitted to prediction errors, the pattern for the two eyespot size traits seems to suggest that directional epistasis might occur for them. Such directional epistasis might simply follow from the shape of threshold traits translating liabilities to traits constrained between 0 and 100%, but the increase of prediction error and thus a small acceleration in the response when EyeP is around 80% argues against that. As we found substantial environmental effect on trait means, we should consider environmental effects on ***G*** too, although it is unclear of which magnitude these are expected to be (Wood & Brodie 2015).

### Conclusion

Our results clearly call for an effort to extend the multivariate breeder’s equation with a suite of mechanistic models that can be fitted to multi-trait data and which allow for environmental effects and different trait-specific mechanisms translating allelic variation into genetic variances, genetic correlations and phenotypes. In our study, the breeder’s equation adequately predicts selection responses, but more mechanistic quantitative genetic models might make it easier to resolve discussions on the adequacy of quantitative genetics by allowing a wider variety of postulated mechanisms to be fitted to data and compared statistically.

## Acknowledgments

We thank K. Koops, N. Wurzer, M. Lavrijsen, and F. Kesbeke for their invaluable assistance with caterpillar and butterfly care, and J. Wolf, W. Frankino, B.J. Zwaan, and P.M Brakefield for discussion and insights at different stages of the project. Comments from M. Blows and D.Emlen greatly improved the manuscript. We thank the Dutch Scientific Organization (NWO) for funding: VENI 863.03.004 (TJMVD), ALW 813.04.002 (supporting CEA), VIDI 864.08.010 (PB).

## Symbols

*G*: Additive genetic variance-covariance matrix
*G*_0_: ***G*** matrix of the starting population (G0)
*P*: ***P*** Phenotypic variance-covariance matrix
***P***_0_: ***P*** matrix of the starting population (G0)
***P***^*^: ***P*** matrix of selected parents only
*s*: Selection differential
*μ*: population mean

## Abbreviations

EyeA: Anterior dorsal forewing eyespot EyeP Posterior dorsal forewing eyespot
Eye4: Fourth eyespot on the ventral hindwing Eye6 Sixth eyespot on the ventral hindwing
Gen0: Starting population for a selection experiment
Gen1- Gen10: Subsequent offspring generations during a selection experiment
A+P+: Concerted selection; both eyespots A, P selected for increased size
A−P−: Concerted selection; both eyespots A, P selected for decreased size
A+P−: Antagonistic selection; A selected for increased and P for decreased size
A−P+: Antagonistic selection; A selected for decreased and P for increased size
4G6G: Concerted selection lines; both eyespots 4 and 6 selected for increased gold
4B6B: Concerted selection lines; both eyespots 4 and 6 selected for increased black
4B6G: Antagonistic selection; 4 selected for increased black, 6 for increased gold
4G6B: Antagonistic selection; 4 selected for increased gold, 6 for increased black
UC: Unselected control lines
ML: Maximum likelihood
AIC: Akaike Information Criterion

